# *C. elegans* avoidance of *Pseudomonas*: thioredoxin shapes the sensory response to bacterially produced nitric oxide

**DOI:** 10.1101/289827

**Authors:** Yingsong Hao, Wenxing Yang, Jing Ren, Qi Hall, Yun Zhang, Joshua M. Kaplan

## Abstract

We show that *C. elegans* avoids a bacterial pathogen *Pseudomonas aeruginosa* (PA14) by detecting PA14-produced nitric oxide (NO). PA14 mutants deficient for NO production fail to elicit avoidance and NO donors repel worms. PA14 and NO avoidance are mediated by the ASJ chemosensory neurons, which respond to NO with intracellular calcium rises. PA14 avoidance and NO-evoked calcium responses require receptor guanylate cyclases (DAF-11 and GCY-27), and cyclic nucleotide gated ion channels (TAX-2 and -4). ASJ exhibits calcium increases at both the onset and removal of NO. These NO-evoked ON and OFF calcium transients are affected by a redox sensing protein, TRX-1/thioredoxin. TRX-1’s trans-nitrosylation activity inhibits the ON transient whereas TRX-1’s de-nitrosylation activity promotes the OFF transient. Thus, *C. elegans* exploits bacterially produced NO as a cue to mediate avoidance and TRX-1 functions as an NO-sensor that endows ASJ with a bi-phasic response to NO exposure.

## Introduction

Nitric oxide (NO) is an important signaling molecule in both prokaryotes and eukaryotes. In mammals, NO regulates key physiological events, such as vasodilation, inflammatory response, and neurotransmission (Feelisch and Martin, 1995). NO regulates innate immunity and life span in the nematode *C. elegans* (Gusarov et al., 2013), as well as virulence and biofilm formation in different bacteria (Cutruzzola and Frankenberg-Dinkel, 2016; Shatalin et al., 2008). NO signaling is mediated by either of two biochemical mechanisms. As a reactive oxygen species, NO covalently modifies the thiol side chain of reactive cysteine residues (forming S-nitrosylated adducts), thereby modulating the activity of these proteins (Foster et al., 2003). NO can also bind to the heme co-factor associated with soluble guanylate cyclases (sGCs), thereby stimulating cGMP production and activating downstream cGMP targets (Denninger and Marletta, 1999).

Almost all living organisms, including bacteria, fungi, plants and animals, are able to produce NO with nitric oxide synthases (NOS) (Ghosh and Salerno, 2003). Due to its small molecular weight and gaseous nature, NO readily diffuses throughout the surrounding tissues to regulate cellular physiology. NO is also released into air, where it may function as an environmental cue. Lightning generates the major abiotic source of environmental NO (Navarro-Gonzalez et al., 2001). Despite its prevalence in the environment, it remains unclear if NO is utilized as a sensory cue by terrestrial animals to elicit behavioral responses. sGCs are the only described sensors for biosynthetically produced NO, mediating NO-evoked muscle relaxation and vasodilation (Gow et al., 2002; Stoll et al., 2001). However, it is unclear if sGCs also play a role in NO-evoked sensory responses. In vertebrates, NO modulates the activities of various ion channels, either directly through S-nitrosylation or indirectly through sGCs. NO regulation of ion channels alters neuron and muscle excitability (Bolotina et al., 1994; Broillet and Firestein, 1996, 1997; Koh et al., 1995; Wang et al., 2012; Wilson and Garthwaite, 2010). For example, in salamander olfactory sensory neurons, S-nitrosylation of a cysteine residue in cyclic nucleotide-gated (CNG) channels activates these channels, thereby directly altering odor-evoked responses in these cells (Broillet and Firestein, 1996, 1997). CNG channels are highly conserved among invertebrates and vertebrates. Because both CNG channels and guanylate cyclases are essential for transducing responses for many sensory modalities, these results suggest that CNG channels and guanylate cyclases may also play a role in NO-evoked sensory responses.

Unlike most metazoans, the nematode *C. elegans* lacks genes encoding NOS (Gusarov et al., 2013) and consequently cannot synthesize NO. Nonetheless, *C. elegans* is exposed to several potential environmental sources of NO, including NO produced by bacteria, which regulates *C. elegans* stress responses and aging (Gusarov et al., 2013). *C. elegans* lives in rotting organic matter, where it feeds on diverse microbes, including the gram-negative bacteria from the *Pseudomonas* and the *Bacillus* genera (Samuel et al., 2016). *C. elegans* exhibits a rich repertoire of behavioral interactions with *Pseudomonas aeruginosa*, *Serratia marascens*, and *Bacillus subtilis* (Brandt and Ringstad, 2015; Garsin et al., 2003; Reddy et al., 2009; Styer et al., 2008). A few bacterially derived metabolites have been shown to mediate these behavioral responses (Brandt and Ringstad, 2015; Meisel and Kim, 2014). Here we test the idea that bacterially produced NO is an ecologically significant environmental cue for *C. elegans-*pathogen interactions.

## Results

### Bacterially derived NO is required for avoidance of *P. aeruginosa* PA14

When cultured with a non-pathogenic *Escherichia coli* strain as the food source, *C elegans* spends most of its time foraging inside the bacterial lawn (Bendesky et al., 2011). By contrast, when feeding on a pathogenic bacterial strain (e.g. *P. aeruginosa* PA14), *C. elegans* avoids the pathogen by foraging off the bacterial lawn (Reddy et al., 2009; Styer et al., 2008). Using a modified assay, we assessed PA14 avoidance by *C. elegans* young adults (detailed in Experimental Procedures). Consistent with prior findings, over a time course of a few hours the majority of the wild-type adult animals remained off the PA14 lawn (Figure 1A). We quantified PA14 avoidance as the percentage of animals inside the PA14 lawn after 8 hours of co-culture (Figure 1B). PA14 avoidance is contingent on the virulence of the bacterial strain, as an isogenic PA14 *gacA* mutant, which is significantly impaired in its ability to kill *C. elegans* (Tan et al., 1999), failed to elicit avoidance of the bacterial lawn (Figure 1C).

**Figure 1.**
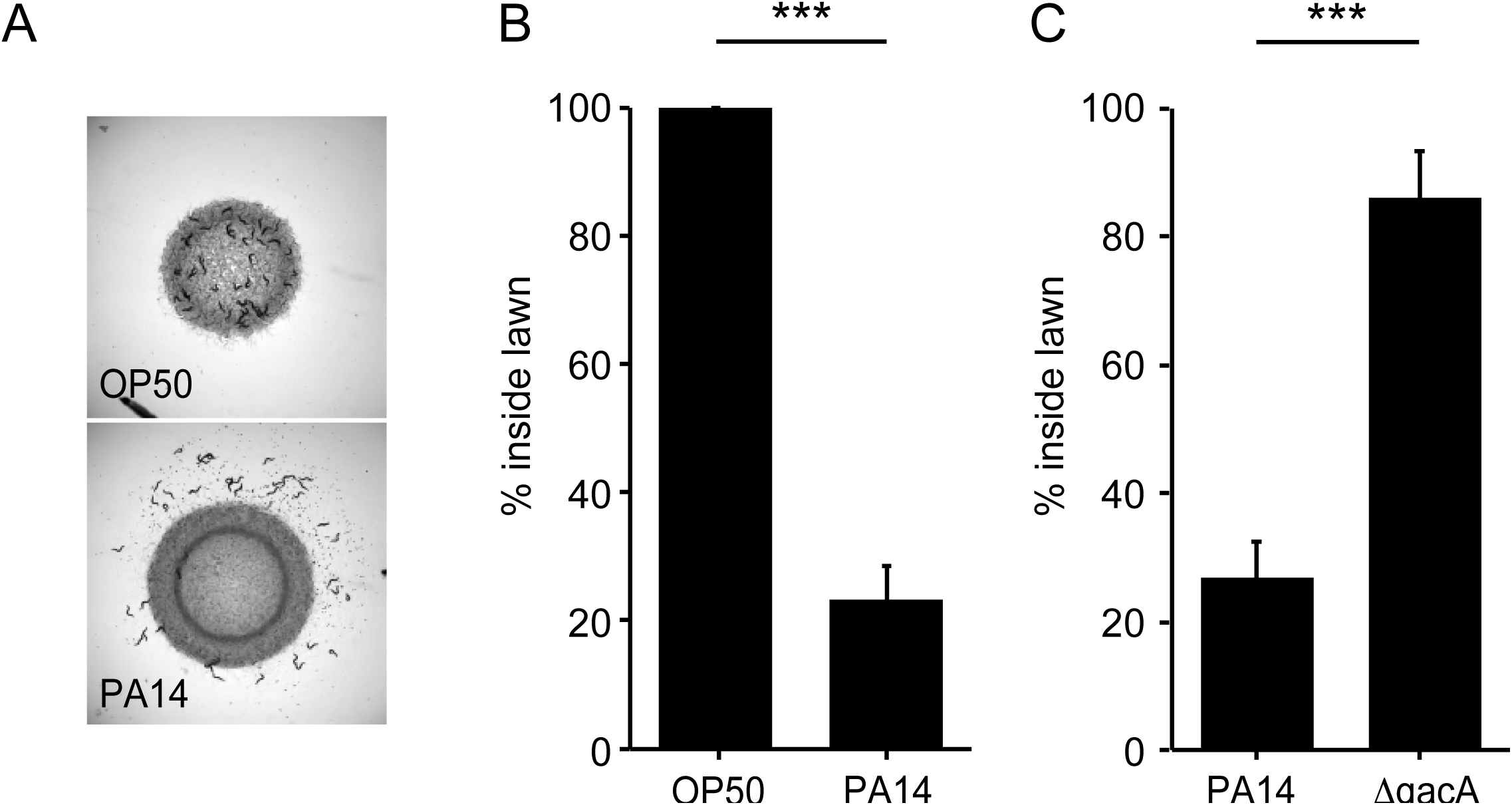
*C. elegans* avoids PA14, but not OP50 or *ΔgacA*. (**A**) Foraging behavior of wild-type animals after 8 hr exposure to *E. coli* OP50 (top) and *P. aeruginosa* PA14 (bottom) lawns is shown. (**B**) Lawn occupancy of wild-type animals on OP50 and PA14 after 8 hr are compared. (**C**) Lawn occupancy of wild-type animals on PA14 vs. *ΔgacA* after 8 hr. ***p < 0.001 as determined by Student’s t-test. Values represent means of four independent experiments. Error bars indicate SEM.

To test the potential role of nitric oxide (NO) in regulating the interaction of *C. elegans* and PA14, we tested a PA14 mutant that was deficient for NO production. *P. aeruginosa* produces NO via a biosynthetic pathway that converts nitrite to NO with nitrite reductase (*nir*) (Figure 2A). Prior studies reported that a PA14 mutant carrying a *nirS* mutation exhibit decreased ability to kill infected *C. elegans*, likely due to the decreased expression of virulence factors (Van Alst et al., 2007; Van Alst et al., 2009). Prompted by these results, we tested the idea that PA14-produced NO elicits avoidance by *C. elegans*. We found that avoidance of *nirS* mutants was dramatically reduced compared to wild-type PA14 controls (Figure 2B). Loss of repulsion by the *nirS* mutant could result from either decreased NO levels or from changes in other virulence factors potentially activated by NO (Van Alst et al., 2007; Van Alst et al., 2009). To distinguish between these possibilities, we asked if chemical NO donors also elicit an avoidance response. When placed inside a non-pathogenic *E. coli* (OP50 strain) lawn, two different NO donors (MAHMA NONOate and DPTA NONOate) elicited *C. elegans* avoidance. After a 30-minute exposure, the majority of animals remained out of the NO-tainted *E. coli* lawn, similarly to the avoidance elicited by the PA14 lawn (Figure 2C). Collectively, these results suggest that bacterially produced NO is required for PA14 avoidance and suggests that *C. elegans* responds to NO as a repulsive chemosensory cue.

**Figure 2.**
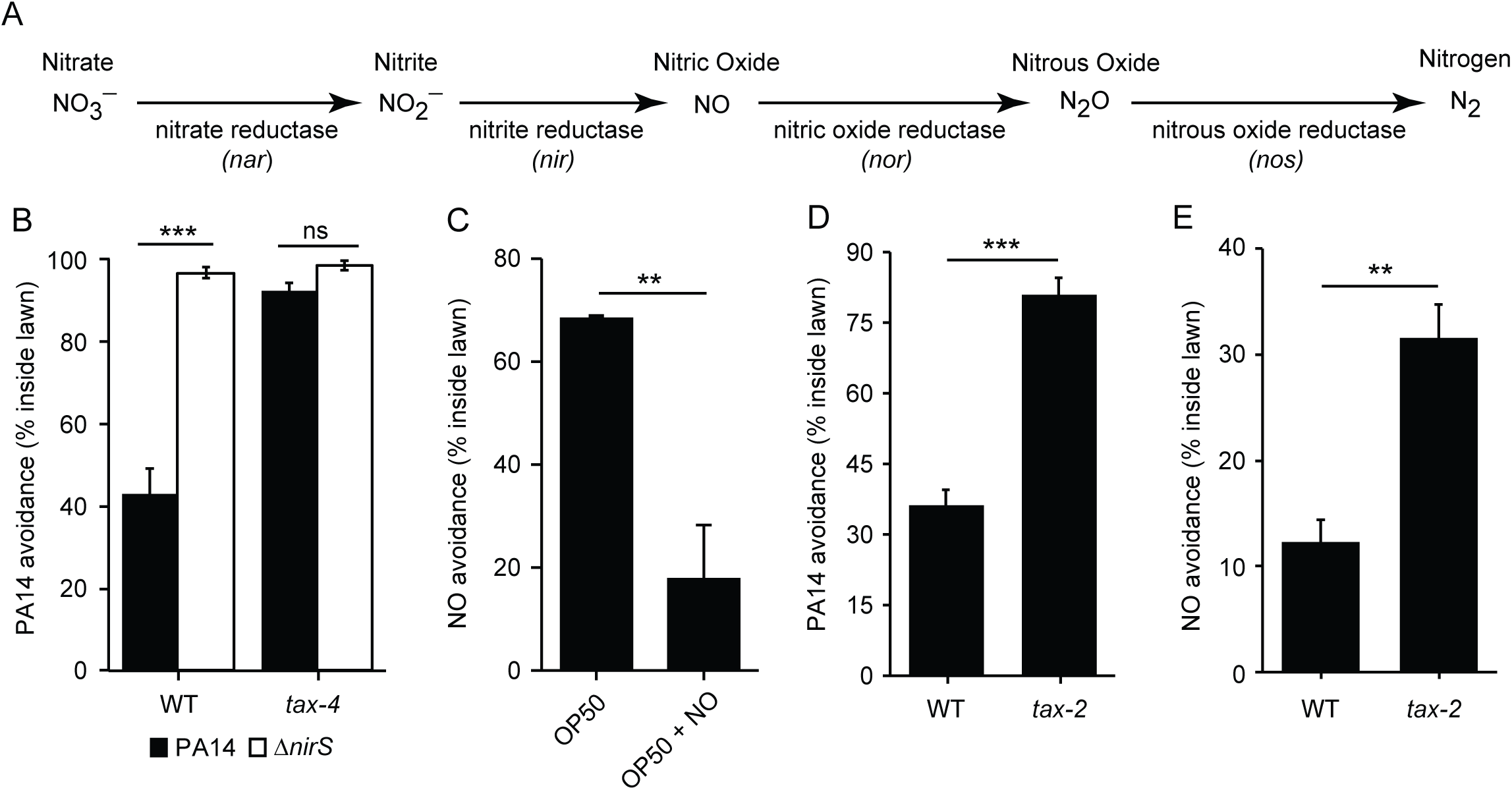
Bacterial NO is required for PA14 avoidance. (**A**) The NO biosynthetic pathway in *P. aeruginosa* is shown. (**B**) Avoidance of PA14 *ΔnirS* lawns was dramatically reduced compared to wild-type PA14. By contrast, neither wild-type nor *nirS* mutants elicited lawn avoidance in *tax-4* mutant worms. (**C**) *E. coli* OP50 (OP50) lawns supplemented with the NO donor DPTA NONOate elicited *C. elegans* avoidance after a 30-minute exposure. (**D-E**) The mutation *tax-2(p671)* decreased PA14 (**D**) and NO donor (**E**) avoidance. **p < 0.01 and ***p < 0.001 as determined by Student’s t-test. Values represent means of four independent experiments. Error bars indicate SEM.

### PA14 and NO avoidance require the TAX-2/TAX-4 CNG channels

Next, we tested the idea that CNG channels are required for PA14 and NO avoidance. CNG channels mediate many *C. elegans* chemosensory responses. The *C. elegans* genome contains several genes that encode CNG channel subunits, including TAX-4/CNGα and TAX-2/CNGβ, which form a heteromeric cGMP-gated cation channel (Komatsu et al., 1999) and are expressed in several classes of chemosensory neurons (Coburn et al., 1998). PA14 and NO donor avoidance was abolished in *tax-4(p678)* mutants (Figure 2B and 3C). Similar PA14 and NO avoidance defects were also observed in the *tax-2(p671)* mutants (Figure 2D and 2E). Thus, TAX-4/CNGα and TAX-2/CNGβ subunits were both required for proper avoidance behavior. Taken together, these results show that the cGMP-gated sensory channel TAX-4/TAX-2 is required for NO and PA14 avoidance. Because TAX-4/TAX-2 channels mediate several chemosensory responses, these results further support the idea that *C. elegans* responds to NO as a repulsive environmental odorant.

**Figure 3.**
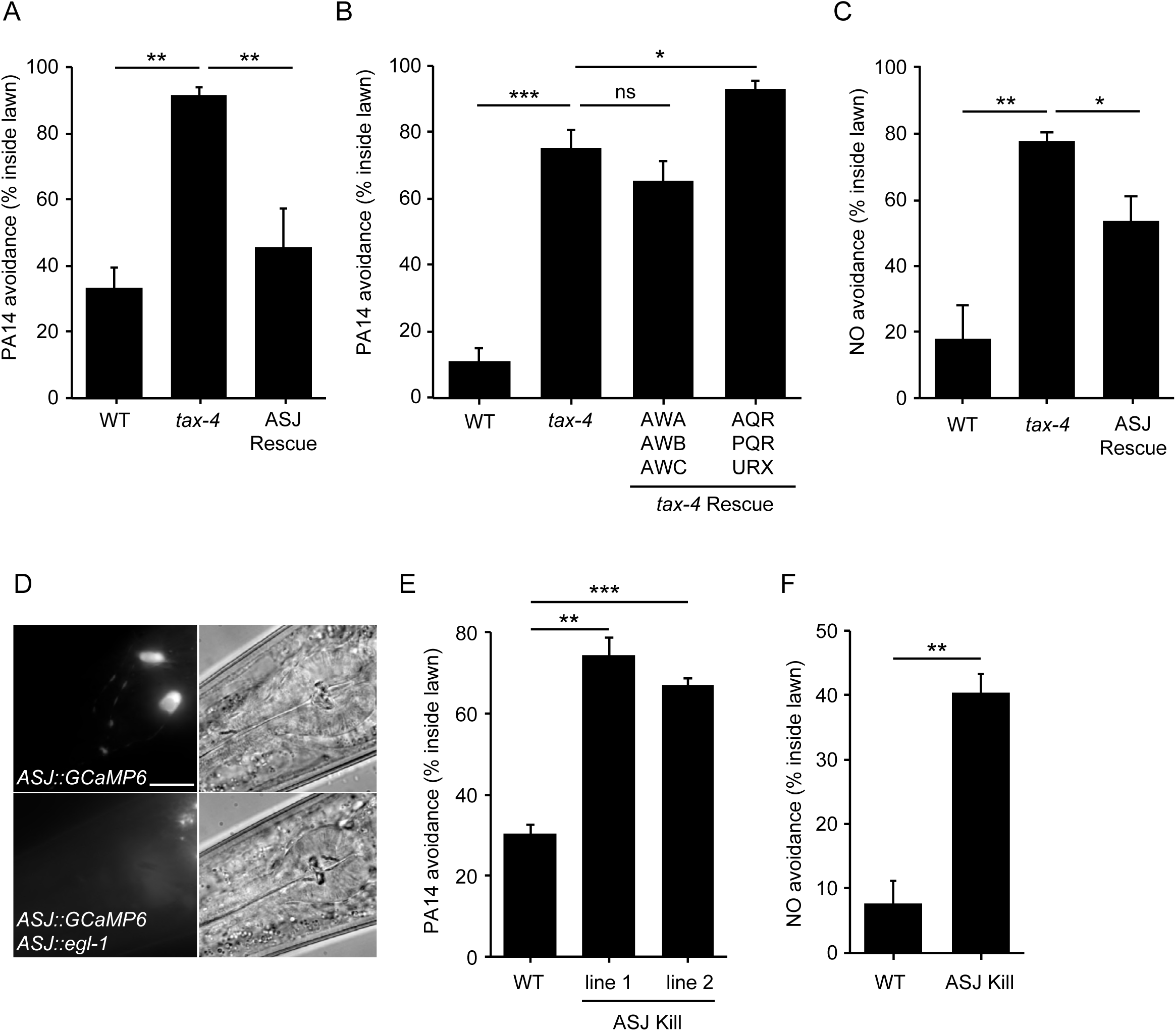
ASJ mediates both PA14 and NO avoidance behavior. (**A, B, C**) The mutation *tax-4(p678)* abolished PA14 (**A, B**) and NO donor (**C**) avoidance. A transgene expressing *tax-4* selectively in ASJ (**A, C**), but not in AWA/AWB/AWC or AQR/PQR/URX (**B**) rescued both PA14 and NO donor avoidance defects. (**D**) Fluorescence (left) and Nomarski (right) images of head region of wild-type animals expressing *ptrx-1::GCaMP6.0* (top), or *ptrx-1::GCaMP6.0* with *pssu-1::egl-1* (bottom). Scale bar indicates 5 μm. (**E, F**) ASJ-ablated animals exhibited significantly decreased PA14 (**E**) and NO donor (**F**) avoidance. *p < 0.05, **p < 0.01, and ***p < 0.001 as determined by Student’s t-test. Values represent means of at least four independent experiments. Error bars indicate SEM. *tax-4* transgenes are as follows: ASJ (*trx-1* promoter), AWA/AWB/AWC (*odr-3* promoter), AQR/PQR/URX (*gcy-36* promoter).

### ASJ neurons mediate NO sensation to elicit avoidance behavior

To identify the sensory neurons mediating PA14 and NO avoidance, we determined which neurons require TAX-4 expression for these responses (Figure 3A-3C). Transgenes expressing a *tax-4* cDNA with either the *odr-3* promoter (expressed in AWA, AWB and AWC olfactory neurons) (Roayaie et al., 1998), or the *gcy-36* promoter (expressed in oxygen sensing AQR, PQR, and URX neurons (Cheung et al., 2004; Gray et al., 2004) both failed to rescue the PA14 avoidance defects of *tax-4* mutants (Figure 3B). By contrast, a transgene expressing *tax-4* selectively in ASJ sensory neurons (using the *trx-1* promoter) rescued both PA14 and NO donor avoidance defects exhibited by *tax-4(p678)* mutants (Figure 3A and 3C). Thus, TAX-4 CNG channels act in the ASJ neurons to promote PA14 and NO donor avoidance. To confirm that ASJ neurons are required for PA14 and NO avoidance, we genetically ablated ASJ neurons by expressing the pro-apoptotic EGL-1 protein (Conradt and Horvitz, 1998) using the *trx-1* promoter (Figure 3D). As expected, inducing ASJ cell death significantly decreased PA14 and NO donor avoidance (Figure 3E-3F). A prior study showed that ASJ neurons respond to two bacterial metabolites to mediate PA14 avoidance (Meisel and Kim, 2014). Here, we show that ASJ neurons also sense PA14-derived NO as a repulsive cue through the TAX-2/TAX-4 CNG channels, thereby promoting avoidance of virulent PA14 strains.

To determine if ASJ neurons are activated by NO, we recorded and quantified intracellular calcium transients in ASJ, using a genetically encoded calcium indicator GCAMP6s (Chen et al., 2013). Worms were confined in a microfluidic device with their nose (and associated chemosensory endings) exposed to fluidic streams of sensory stimuli delivered with precise temporal control (Chronis et al., 2007). Exposure to NO donor evoked a significant increase in GCaMP6s fluorescence in the ASJ neurons, reaching peak intensity within a few seconds and gradually returning to baseline fluorescence despite the continuing exposure to NO (Figure 4A). Removing the NO stimulus also evoked an ASJ calcium transient that lasted a few seconds (Figure 4A). By contrast, switching between streams of control buffer solution did not alter the GCaMP6s signal in ASJ (Figure 4B). To determine if ASJ neurons sense NO directly, we examined *unc-13(s69)* mutants, in which synaptic transmission is nearly completely blocked (Richmond et al., 1999). We found that the *unc-13* mutation did not alter either the ON or OFF responses of ASJ to NO (Figure 4C), suggesting that NO-evoked ASJ calcium transients are unlikely to result from indirect activation of ASJ by synaptic input. Collectively, these results indicate that the ASJ sensory neurons directly sense NO, and that ASJ neurons have a biphasic response to NO, exhibiting increased cytoplasmic calcium at both NO onset and removal (hereafter indicated as ON and OFF responses).

**Figure 4.**
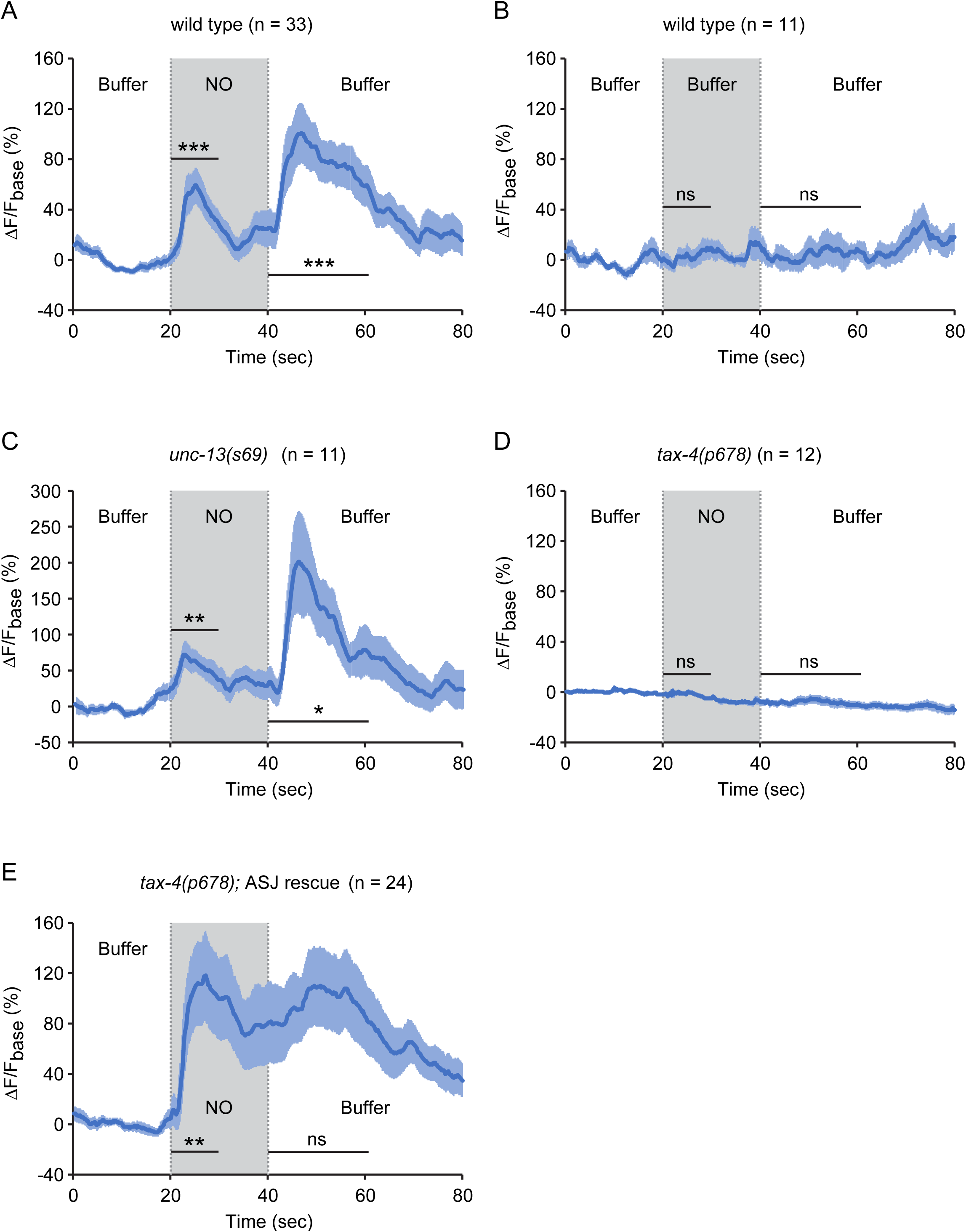
The sensory neurons ASJ respond to NO. (**A, B**) ASJ neurons respond to the onset and removal of NO with increased GCaMP6 signal (**A**) but do not respond when switched between control buffer (**B**). (**C**) Blocking synaptic transmission (in *unc-13* mutants) had little effect on ASJ responses to NO. (**D, E**) The *tax-4(p678)* mutation abolished the ASJ response to the onset and the removal of NO (**D**). Expressing a wild-type *tax-4* cDNA in ASJ rescued ASJ response to the onset of NO stimulation (**E**). ΔF = F-F_base_, F_base_ is the average intensity over the 20 seconds of the recording before stimulus onset, solid line represents mean and shade represents SEM, average ΔF/F_base_% for the 3-s window before NO onset is compared with average ΔF/F_base_% for the 10-s window after the onset ((Wilcoxon signed rank test for data that were not normally distributed (A, E); paired Student’s *t-*test for normally distributed data (B-D)); average ΔF/F_base_% for the 3-s window before NO removal is compared with average ΔF/F_base_% for the 20-s window after the removal ((Wilcoxon signed rank test for data that were not normally distributed (A, B, D, E); paired Student’s *t-*test for normally distributed data (C)), *** p < 0.001, ** p < 0.01, * p < 0.05, ns: not significant (Materials and Methods).

To determine if CNG channels are required for NO-activation of ASJ, we recorded the ASJ GCaMP6s signal in *tax-4* mutants. The *tax-4(p678)* mutation, which abolished PA14 and NO donor avoidance (Figure 2B, 3A-C), also eliminated the NO-evoked ON and OFF calcium transients in ASJ (Figure 4D). A transgene selectively restoring *tax-4* expression in ASJ reinstated the NO-evoked ON calcium transients in ASJ (Figure 4E). These results indicate that activation of TAX-4 channels in the ASJ sensory neurons generates increased calcium transients in response to NO exposure, which results in NO avoidance behavior.

### The guanylate cyclase DAF-11 mediates NO sensation in ASJ

The requirement for TAX-2/TAX-4 CNG channels for NO sensation suggests that NO stimulates cGMP synthesis in ASJ. Responses to extracellular gaseous ligands, such as O_2_ and NO, are often mediated by sGCs; however, none of the sGCs encoding genes is expressed in ASJ neurons (www.wormbase.org). Receptor guanylate cyclases (rGCs) mediate *C. elegans* responses to CO_2_ (Hallem et al., 2011). Prompted by these results, we tested the idea that rGCs mediate NO responses. ASJ neurons express two rGCs (GCY-27 and DAF-11). We found that PA14 and NO donor avoidance were both abolished in *daf-11(m47)* mutants and both avoidance responses were reinstated by a transgene that expresses *daf-11* in the ASJ neurons (Figure 5A). Next, we analyzed ASJ GCaMP6s signals in *daf-11* mutants. We found that the *daf-11(m47)* mutation completely abolished the ASJ ON and OFF response to NO donor (Figure 5C). Expressing the *daf-11* cDNA specifically in ASJ partially rescued the ASJ ON response to NO donor (Figure 5D). In contrast to DAF-11, mutations inactivating GCY-27 decreased but did not abolish PA14 avoidance and had no effect on the ASJ ON response, although they did eliminate the ASJ OFF response to NO donor (Figure 5B and 5E). Together, these results indicate that the rGCs DAF-11 and GCY-27 function in the ASJ neurons to mediate NO sensation. Next, we examined how DAF-11 mediated the sensory response to NO. DAF-11 does not contain a heme-NO-binding domain, although it contains a heme-NO-binding associated (HNOBA) domain. To test its functional importance, we used CRISPR to delete the *daf-11* HNOBA domain. The resulting *daf-11(nu629)* mutation abolished the ASJ response to both the onset and the removal of NO (Figure 5F), suggesting that DAF-11 mediates NO sensing by interacting with other NO-binding proteins.

**Figure 5.**
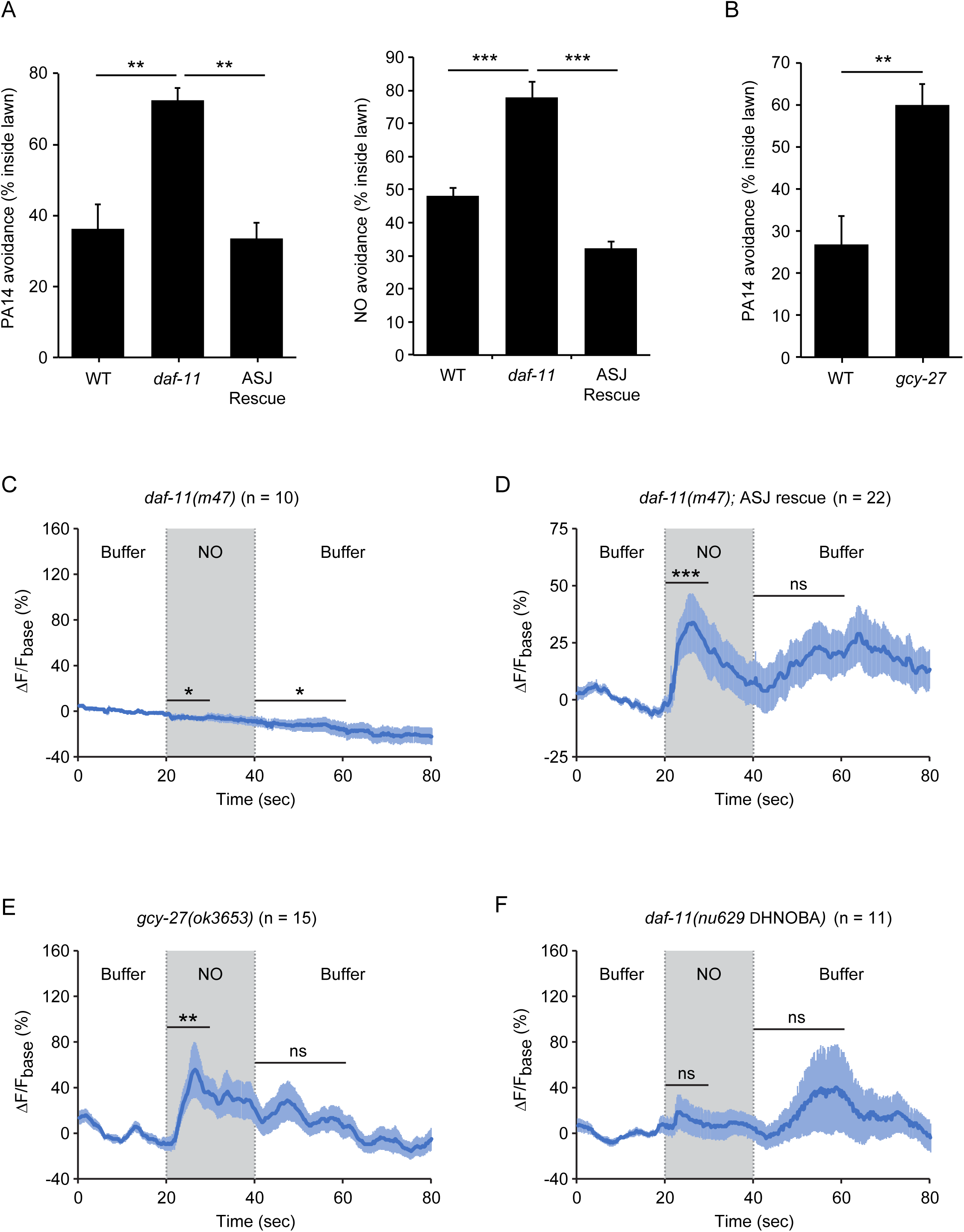
The rGCs DAF-11 and GCY-27 are required for ASJ responses to NO. (A) The *daf-11(m47)* mutation abolished PA14 (left) and NO donor (right) avoidance, both of which were reinstated by a transgene that expresses *daf-11* in the ASJ neurons. (B) The *gcy-27(ok3653)* mutation decreased PA14 avoidance. For **A-B**, **p < 0.01, ***p < 0.001, as determined by Student’s t-test. Values represent means of four independent experiments. Error bars indicate SEM. (**C, D**) The *daf-11(m47)* mutation abolished the NO-evoked calcium transients in the ASJ neurons (**C**). Expressing a wild-type *daf-11* cDNA in ASJ rescued the NO-evoked ON response (**D**). Note, NO-onset and removal slightly suppressed the ASJ GCaMP6 signal in *daf-11(m47)* mutants. (E) The *gcy-27(ok3653)* mutation eliminated the ASJ OFF response to NO. (F) The *daf-11(nu629* ΔHNOBA) mutation abolished the ASJ ON and OFF responses to NO. For **C-F**, ΔF = F-F_base_, F_base_ is the average intensity over the 20 seconds of the recording before stimulus onset, solid line represents mean and shade represents SEM, average ΔF/F_base_% for the 3-s window before NO onset is compared with average ΔF/F_base_% for the 10-s window after the onset ((Wilcoxon signed rank test for data that were not normally distributed (D-F); paired Student’s *t-*test for normally distributed data (C)); average ΔF/F_base_% for the 3-s window before NO removal is compared with average ΔF/F_base_% for the 20-s window after the removal ((Wilcoxon signed rank test for data that were not normally distributed (D, F); paired Student’s *t-* test for normally distributed data (C, E)), *** p < 0.001, ** p < 0.01, * p < 0.05, ns: not significant (Materials and Methods).

### NO acts as an external sensory cue to elicit avoidance

NO is freely diffusible and membrane permeable; consequently, PA14 produced NO could act as either an external sensory cue or by directly activating intracellular signaling pathways in ASJ. We did several experiments to distinguish between these possibilities. If NO acts as an external chemosensory cue, PA14 and NO donor avoidance should be diminished in mutants that have defective ciliated sensory endings. Previous studies identified mutations in genes encoding the components of ciliated sensory endings that disrupt the function of chemosensory neurons, including the ASJ neurons (Perkins et al., 1986). Two cilia defective mutants, *osm-12(n1606)* and *bbs-9(gk471)*, were both defective in avoiding the lawn of PA14 and the NO donor (Figure 6A and 6B). These results indicate that the normal function of the sensory cilia is required for the NO sensation.

**Figure 6.**
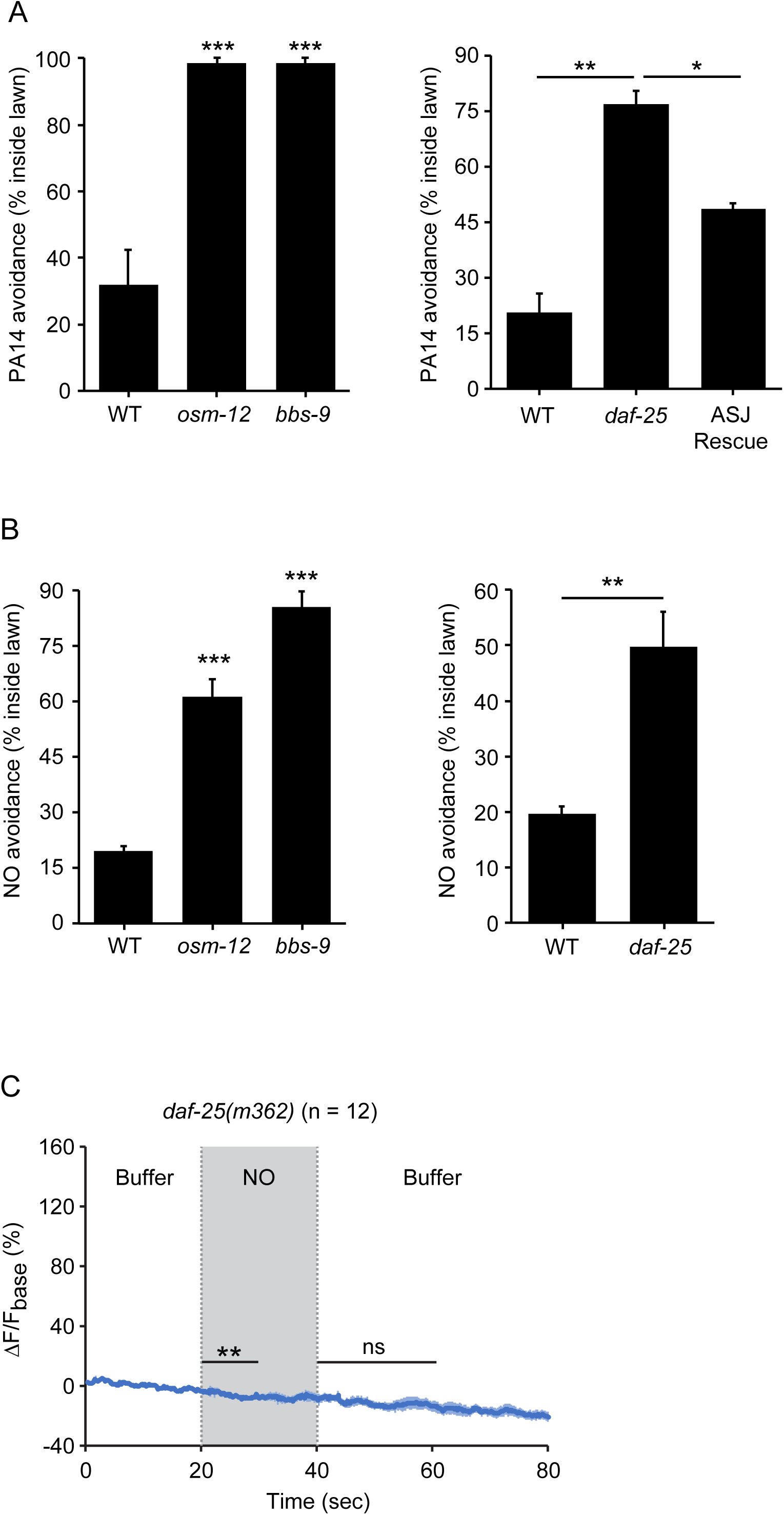
*C. elegans* senses NO as an external cue. (**A, B**) *osm-12(n1606), bbs-9(gk471) and daf-25(m362)* mutants were defective for PA14 (**A**) and NO donor (**B**) avoidance. Expressing a wild-type *daf-25* cDNA in ASJ partially rescued the *daf-25* mutant PA14 avoidance defect (**A**). *p < 0.05, **p < 0.01, and ***p < 0.001 as determined by Student’s t-test. Values represent means of four independent experiments. Error bars indicate SEM. The *daf-25* ASJ transgene was expressed by the *ssu-1* promoter. (**C**) *daf-25(m362)* mutants lacked ASJ responses to the onset and removal of NO. ΔF = F-F_base_, F_base_ is the average intensity over the 20 seconds of the recording before stimulus onset, solid line represents mean and shade represents SEM, average ΔF/F_base_% for the 3-s window before NO onset is compared with average ΔF/F_base_% for the 10-s window after the onset; average ΔF/F_base_% for the 3-s window before NO removal is compared with average ΔF/F_base_% for the 20-s window after the removal, paired Student’s *t-*test for normally distributed data, ** p < 0.01, ns: not significant (Materials and Methods). NO-onset slightly suppressed the GCaMP6 signal in ASJ.

To further address this question, we asked if DAF-11 rGC and TAX-2/4 CNGs must be localized to ASJ sensory endings to mediate NO responses. To test this idea, we analyzed *daf-25(m362)* mutants, which lack a cargo adaptor that is required for rGC and CNG transport to ciliated sensory endings (Fujiwara et al., 2010; Jensen et al., 2010; Wojtyniak et al., 2013). Avoidance of NO and the PA14 lawn were both defective in *daf-25(m362)* mutants (Figure 6A and 6B). The defect in PA14 avoidance was partially rescued by a transgene that restores DAF-25 expression in ASJ neurons (Figure 6A). As in *daf-11(m47)* mutants, the NO evoked ON and OFF calcium transients in ASJ were completely abolished in *daf-25(m362)* mutants (Figure 6C). Thus, the ability of ASJ neurons to detect NO requires DAF-11 and TAX-2/4 CNG channel localization to ciliated sensory endings. Taken together, these results show that NO is sensed by ASJ as an external sensory odor.

### TRX-1/Thioredoxin shapes the ASJ response to NO

How do ASJ neurons detect NO? NO covalently modifies reactive cysteine residues by S-nitrosylation (SNO). To determine if protein-SNO modifications are involved in NO detection, we analyzed mutants lacking protein de-nitrosylating enzymes. Protein-SNO modifications are reversed by two classes of enzymes, Thioredoxins (TRX) and nitrosoglutathione reductases (GSNOR) (Benhar et al., 2009); consequently, protein-SNO adducts should accumulate in mutants lacking TRX and GSNOR. The *C. elegans* genome encodes multiple thioredoxin genes. We focused on the *trx-1* gene (Figure 7A) because it is exclusively expressed in the ASJ neurons (Miranda-Vizuete et al., 2006). The amplitude and duration of the NO-evoked ON transient in ASJ were both significantly increased in *trx-1(jh127)* null mutants (Figure 7A and 7B) and this defect was rescued by a single copy transgene restoring TRX-1 expression in ASJ neurons (Figure 7C). To determine the effect of de-nitrosylation in NO sensing, we used CRISPR to isolate a catalytically inactive *trx-1(nu517)* mutant, containing the C38S mutation in the active site for de-nitrosylation (Figure 7A). In *trx-1(nu517* C38S) mutants, the NO evoked OFF calcium transient was eliminated, whereas the ON transient was unaltered (Figure 7D). These results suggest that TRX-1’s de-nitrosylating activity is required for ASJ to generate increased cytoplasmic calcium in response to removing NO. Interestingly, the ASJ NO donor responses of *trx-1(jh127)* null mutants differed significantly from those in *trx-1(nu517* C38S) mutants (compare Figures 7B and 7D), suggesting that the null phenotype cannot be explained by decreased de-nitrosylation activity. Cysteine residues outside of the catalytic domain are thought to promote other TRX functions. For example, TRX promotes nitrosylation of other proteins, and this trans-nitrosylation activity is eliminated by mutations altering Cys-72 (Mitchell and Marletta, 2005). To address the role of trans-nitrosylation, we rescued *trx-1* null mutants with a transgene expressing TRX-1(C72S) (Figure 7A). The ASJ NO donor responses observed in TRX-1(C72S) transgenics were similar to those found in *trx-1* null mutants (Figure 7E). These results suggest that the exaggerated amplitude and prolonged time course of the ASJ ON response to NO donor results from inactivation of TRX-1’s trans-nitrosylation activity.

**Figure 7.**
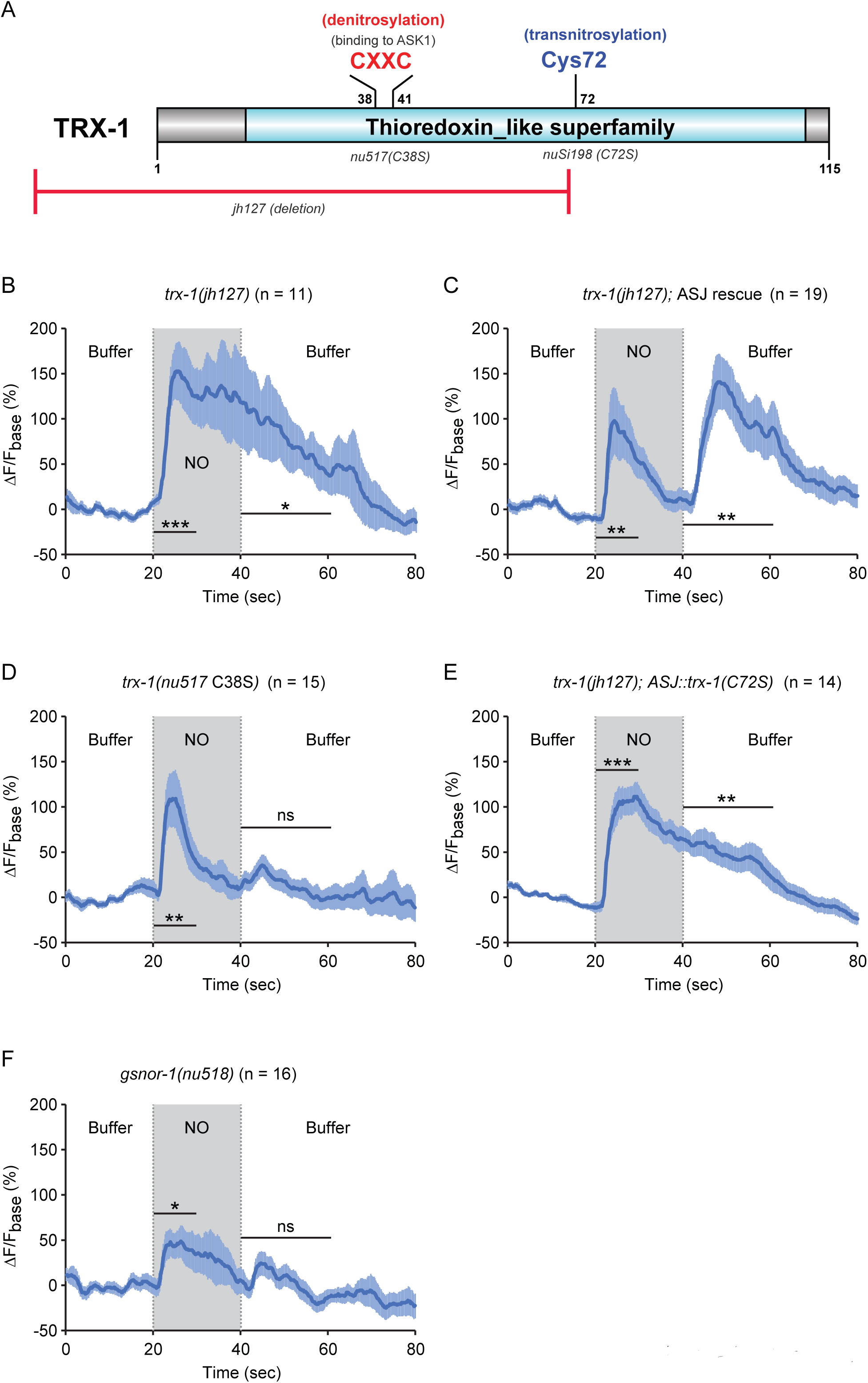
TRX-1/Thioredoxin regulates the ASJ response to NO. (**A**) Domain organization of *C. elegans* thioredoxin TRX-1. Cysteines forming the de-nitrosylation active site (Cys38XXCys41) in TRX-1 are highlighted in red. Cys72, which is involved in trans-nitrosylation is highlighted in blue. The deleted region in *jh127* as well as point mutations in *nu517* (C38S) and *nuSi198* (C72S) are indicated. (**B, C**) A *trx-1(jh127)* null mutant exhibited a long-lasting ASJ response to the onset of NO simulation, which slowly returned to baseline following NO removal (**B**). This defect was rescued by a single copy transgene expressing wild-type *trx-1* in ASJ (**C**). (D) A mutation in the active site for de-nitrosylation, *trx-1(nu517* C38S) eliminated the ASJ response to NO removal. (E) In contrast to the full rescue by wild type TRX-1 (**C**), expressing a mutant TRX-1(C72S) single copy transgene failed to rescue the defective ASJ response to NO in *trx-1(jh127)* null mutants. (F) A mutation inactivating another de-nitrosylation enzyme (*gsnor-1*) disrupted the response of ASJ to the removal of NO stimulation, similar to the phenotype exhibited by the de-nitrosylation defective *trx-1(nu517* C38S) mutant (**D**). ΔF = F-F_base_, F_base_ is the average intensity over the 20 seconds of the recording before stimulus onset, solid line represents mean and shade represents SEM, average ΔF/F_base_% for the 3-s window before NO onset is compared with average ΔF/F_base_% for the 10-s window after the onset ((Wilcoxon signed rank test for data that were not normally distributed (C); paired Student’s *t-*test for normally distributed data (B, D-F)); average ΔF/F_base_% for the 3-s window before NO removal is compared with average ΔF/F_base_% for the 20-s window after the removal ((Wilcoxon signed rank test for data that were not normally distributed (B, C); paired Student’s *t-* test for normally distributed data (D-F)), *** p < 0.001, ** p < 0.01, * p < 0.05, ns: not significant (Materials and Methods).

To further investigate the role of protein-SNO modifications in ASJ responses to NO donors, we analyzed mutants lacking a second de-nitrosylating enzyme GSNOR. The *C. elegans* genome encodes a single GSNOR gene, H24K24.3 (hereafter designated *gsnor-1*). We used CRISPR to isolate an early nonsense mutation in *gsnor-1(nu518)*. The NO evoked ASJ OFF response was significantly diminished in the *gsnor-1* mutants, whereas the ON response remained (Figure 7F). Thus, analysis of *trx-1* and *gsnor-1* mutants both suggest that protein de-nitrosylation is required for the ASJ response evoked by removing NO.

## Discussion

Here we show that bacterially produced NO elicits *C. elegans* avoidance of pathogenic PA14. This avoidance response is mediated by a specific chemosensory neuron (ASJ) and requires NO-mediated activation of receptor GCs and cGMP gated ion channels. Below we discuss the significance of these findings.

### NO as an environmental cue

*C. elegans* is among only a few organisms that do not synthesize NO and, likely, acquires NO from the environment. We show that NO is sensed by a chemosensory neuron (ASJ), NO sensing requires functional ciliated sensory endings, and that NO sensing is defective in *daf-25* mutants (which lack DAF-11 and TAX-4 localization to cilia) (Fujiwara et al., 2010; Jensen et al., 2010; Wojtyniak et al., 2013). Collectively, these results indicate the role of NO as an external sensory cue that represents environmental information to worms.

Several results suggest that ASJ neurons are the primary NO-sensing neurons. NO-avoidance is strongly defective following genetic ablation of ASJ neurons. Similarly, *tax-4* CNG and *daf-11* rGC mutant defects in NO avoidance are rescued by transgenes restoring expression of these genes selectively to ASJ neurons. Despite these results, it remains possible that other neurons also contribute to NO-evoked behaviors. Seven heme-containing sGCs (*gcy-31-37*) are expressed in oxygen sensing neurons AQR, PQR, URX, and BAG (Cheung et al., 2004; Gray et al., 2004; Zimmer et al., 2009). Several sGCs, including GCY-35, mediate sensory responses to oxygen but also display a low affinity binding to NO (Gray et al., 2004). Thus, these oxygen sensing neurons may also respond directly to NO.

### rGCs and CNG channels are required for NO sensing

The only previously described NO sensors are sGCs that bind NO via their associated Heme co-factor (Malinski and Taha, 1992). Here we show that NO-sensing by ASJ neurons is mediated by two rGCs (DAF-11 and GCY-27). Interestingly, a prior study showed that another rGC (GCY-9) mediates CO_2_ detection by the BAG neurons (Hallem et al., 2011). Thus, rGCs mediate detection of two environmental gasses (NO and CO_2_) by *C. elegans*. In addition, we show that NO-sensing requires the function of the cyclic nucleotide-gate (CNG) channel subunit TAX-4 in the ASJ neurons. Together, our results identify rGCs and CNG channels as one underlying mechanism for NO sensation.

How does *daf-11* mediate NO responses? While DAF-11 is required for both the onset and removal response to NO, neither DAF-11 nor GCY-27 contains a Heme-NO-binding (HNOB) domain. Instead, DAF-11 contains a Heme-NO-binding associated (HNOBA) domain, suggesting that DAF-11 mediates NO sensing by interacting with other NO-binding proteins. Consistent with this possibility, removing the HNOBA domain from DAF-11 abolished the ASJ calcium response to NO. In addition to NO sensing, *daf-11* mediates chemosensory responses to several volatile chemicals (Birnby et al., 2000). DAF-11 may regulate sensory responses to different cues by acting together with different signaling molecules. The complete transcriptome of ASJ is not yet available, which would aid the characterization of other factors that regulate various sensory transduction pathways in ASJ.

How do DAF-11 and GCY-27 detect NO? NO could be detected by S-nitrosylation of cysteine residues in DAF-11 and GCY-27 (or proteins associated with them), thereby activating the GC catalytic domain. Consistent with this idea, mutations inactivating the de-nitrosylating enzymes TRX-1/Thioredoxin and GSNOR-1 eliminate the NO-evoked OFF transient in ASJ, while having little effect on the ON transient. Thus, accumulation of protein SNO-adducts or SNO-Glutathione adducts (in the de-nitrosylation defective mutants) was associated with a loss of the ASJ response to removing NO. Alternatively, the transmembrane GCs may associate with other NO binding proteins, such as Heme-containing globins. In this regard, it is interesting that DAF-11, like the mammalian atrial natriuretic peptide (ANP) receptors, contains a conserved cytoplasmic HNOBA domain, which is similar to PAS domains. HNOBA/PAS domains are thought to be directly bound by HSP90, a chaperone that catalyzes incorporation of heme groups into sGCs (Ghosh and Stuehr, 2012; Sarkar et al., 2015). Interestingly, mutations in *daf-21*/HSP90 mimic all of the phenotypes found in *daf-11* mutants, indicating that DAF-11 function requires its interaction with HSP90 (potentially through binding to DAF-11’s HNOBA/PAS domain) (Birnby et al., 2000). Consistent with this possibility, deleting the DAF-11 HNOBA domain abolished the NO response in ASJ. Thus, the coordinated action of DAF-21/HSP90 and DAF-11 in NO sensing could indicate that DAF-11 associates with other heme-binding proteins (e.g. globins). GCY-27 lacks the HNOBA/PAS domain, and consequently would have to detect NO by a distinct mechanism.

Our results suggest that DAF-11 and GCY-27 mediate different aspects of the NO response. GCY-27 is required for the NO-evoked OFF transient whereas DAF-11 is required for both the ON and OFF transients. The mechanism underlying this difference is not known but could reflect a differential GC activation by increasing (DAF-11) and decreasing (DAF-11 and GCY-27) NO concentration. A similar mechanism was previously proposed for detecting increasing (GCY-35/36) and decreasing (GCY-31/33) O_2_ concentrations by distinct cytoplasmic GCs (Zimmer et al., 2009). In this case, we predict that co-expression of DAF-11 and GCY-27 allows ASJ to detect both increasing and decreasing NO concentration, thereby producing ON and OFF transients. Because NO and PA14 avoidance are deficient in both *daf-11* and *gcy-27* mutants, our results also suggest that ON and OFF transients are both required to promote avoidance behavior.

### TRX-1 shapes ASJ’s bi-phasic response to NO

TRX proteins are redox sensitive proteins that have been proposed to play several important roles in cellular responses to NO. TRX has an enzymatic activity that removes SNO-protein and SNO-glutathione adducts (Benhar et al., 2009). This de-nitrosylation activity is mediated by a pair of active site cysteine residues (C38 and C41 in TRX-1). Thioredoxins have also been proposed to promote nitrosylation of other protein substrates, and this trans-nitrosylation activity requires a third cysteine residue (C72 in TRX-1) (Mitchell and Marletta, 2005). Our results suggest that TRX-1 regulates the NO-evoked ASJ response via two distinct activities. TRX-1 inhibits and shortens the NO-evoked ON response, as indicated by a larger and more prolonged ON response in *trx-1* null mutants. This inhibitory function of TRX-1 is eliminated in the C72S mutant, implying that inhibition is mediated by the TRX-1’s trans-nitrosylation activity (Mitchell and Marletta, 2005). Two results suggest that TRX-1’s de-nitrosylating activity promotes the NO-evoked OFF response in ASJ. The OFF response was eliminated in both *trx-1* mutants containing a mutation in the de-nitrosylation active site (*nu517* C38S) and in *gsnor-1* mutants (which lack a second de-nitrosylating enzyme) (Benhar et al., 2009).

Collectively, our results suggest that TRX-1 acts as an NO-sensor that shapes ASJ’s bi-phasic response to NO (Figure 8). Specifically, we propose that during NO exposure TRX-1’s active site cysteines (C38/41) are oxidized, thereby decreasing de-nitrosylation (Engelman et al., 2016; Wang et al., 2014). Oxidation of C38 and 41 promotes C72 nitrosylation (Barglow et al., 2011), which increases trans-nitrosylation activity. Thus, during NO exposure the net effect of TRX-1 would be increased trans-nitrosylation of protein substrates. We propose that these trans-nitrosylated proteins inhibit the ON response. Following NO-removal, TRX-1 active site cysteines are reduced, thereby enhancing TRX-1’s de-nitrosylation activity and inhibiting its trans-nitrosylation activity. As a result, following NO removal, inhibitory SNO-protein adducts formed during NO exposure could be removed by TRX-1’s de-nitrosylating activity, giving rise to the OFF response. Thus, the combined activities of TRX-1 produce ASJ’s bi-phasic ON and OFF responses to NO.

**Figure 8.**
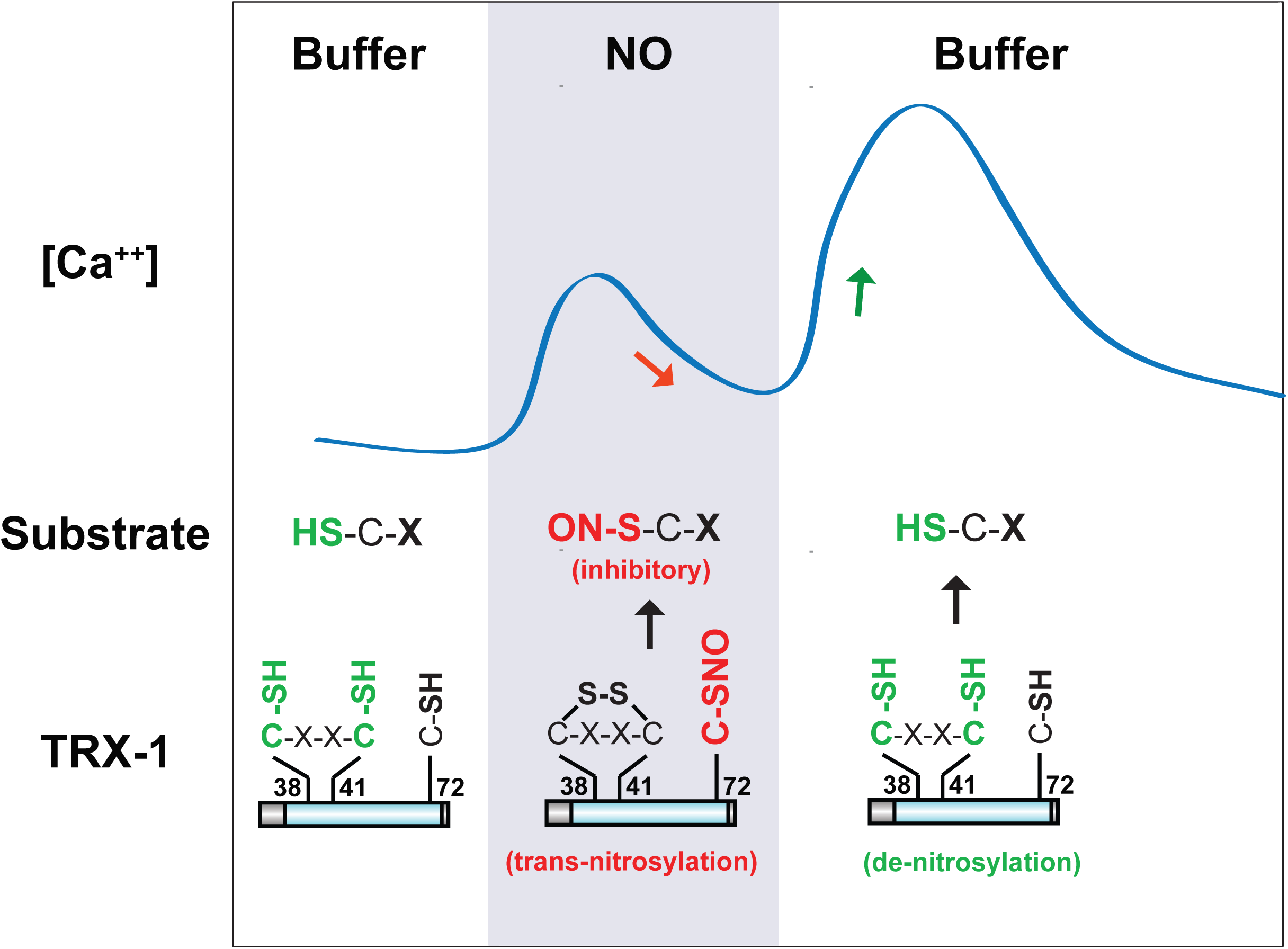
A model for TRX-1/Thioredoxin function in NO sensation. Our data suggest that two enzymatic functions of TRX-1 endow ASJ neurons with a bi-phasic response to environmental NO. The trans-nitrosylation activity is proposed to be active during NO exposures. Trans-nitrosylation of an unidentified protein substrate is proposed to inhibit the ASJ ON response. The de-nitrosylation activity is proposed to be active following NO removal and is proposed to activate the ASJ OFF response (by reversing the inhibitory SNO-protein adducts formed during NO exposures). This model is described in greater detail in the Discussion section.

### Metabolite sensing in pathogen avoidance

A prior study suggested that bacterially produced CO_2_ was utilized as a contextual cue to promote *C. elegans* avoidance of another pathogen, *Serratia marascens* (Brandt and Ringstad, 2015). These authors proposed that environmental CO_2_ indicates proximity to metabolically active bacteria, and that pathogen avoidance is mediated by the coincident detection of CO_2_ in conjunction with other virulence factors. Such a coincidence detection strategy is proposed to optimize the ability of *C. elegans* to forage for nutritional resources since it would allow worms to actively feed on avirulent bacteria while avoiding ingestion of pathogenic bacteria. Our results suggest that *C. elegans* avoidance of PA14 is mediated by a similar coincidence detection strategy. In particular, we propose that recent exposure to NO enhances the repulsive effects of other PA14 metabolites, e.g. phenazine compounds (Meisel and Kim, 2014). Thus, our results combined with this prior study suggest that *C. elegans* discriminates between benign and pathogenic microbes by coincident detection of multiple bacterial metabolites, including both CO_2_ and NO.

## Materials and methods

### C. elegans strains

Strains were maintained at 20°C as described (Brenner, 1974). The wild-type reference strain was N2 Bristol. The mutant strains used were: LGI, *tax-2(p671), unc-13(s69), bbs-9(gk471), daf-25(m362);* LGIII, *tax-4(p678), osm-12(n1606);* LGIV, *gcy-27(ok3653);* LGV, *daf-11(m47), daf-11(ks67).* Descriptions of allele lesions can be found at http://www.wormbase.org.

CRISPR gene editing was utilized to isolate *daf-11(nu629* ΔHNOBA), *trx-1(nu517 C38S)*, and *gsnor-1(nu518)* mutations, using *dpy-10* as a co-CRISPR marker (Ward, 2015). The *gsnor-1* gene corresponds to the H24K24.3 gene in wormbase. Predicted amino acid sequence of these alleles are as follows (mutant residues underlined, * indicates stop codons):

*daf-11(nu629* ΔHNOBA): TQGLNETVKN**<u>EVGRI</u>**ELLPKSVANDLKN

*trx-1(nu517 C38S):* EKIIILDFYATW**<u>S</u>**GPCKAIAPLYKE

*gsnor-1(nu518):* KTNLCQKIRI**<u>**</u>**GNGFMPDGSSRFTCNG

### Constructs and transgenes

All plasmids were derivatives of pPD49.26 (Fire, 1997) and constructed utilizing standard methods. A 1028bp *trx-1* promoter and a 543bp *ssu-1* promoter were used for expression in ASJ. A 3kb *odr-3* promoter was used for expression in AWA/AWB/AWC. The 1089bp *gcy-36* promoter and the *tax-4* cDNA were derived from pEM4 (Gift from Cori Bargmann). GCaMP6.0s was a gift from Jihong Bai. The cDNAs of *egl-1*, *daf-11* and *daf-25* were cloned from a cDNA library using primers corresponding to the predicted start and stop codons of each gene. Full descriptions of all plasmids are available upon request.

Transgenic animals were generated by injecting wild type or mutants with the transgene (10∼25ng/ul) mixed with the co-injection marker, KP#1480 (*pmyo-2::NLS-mCherry*, 10ng/μl), using standard methods (Mello et al., 1991). *nuIs556* was generated by injecting wild type with KP#3311*(ptrx-1::GCaMP6.0s)* at 25ng/ul mixed with the co-injection marker KP#2186 (*punc-122::mCherry)* at 70ng/ul, followed by UV irradiation. The single copy transgenes *nuSi196* [*pssu-1::daf-11*], *nuSi197* [*ptrx-1::trx-1*], and *nuSi198* [*ptrx-1::trx-1(C72S)*] were isolated by the miniMos method (Frokjaer-Jensen et al., 2014).

### Behavior assays

#### The avoidance of the PA14 lawn and full-lawn killing assay

Plates for avoidance assays were prepared as previously described (Tan et al., 1999). An overnight culture of OP50, PA14, PA14 *ΔgacA* or PA14 *nirS* mutant was grown in 5 ml Luria broth (LB) at 37°C. 10 ul of the culture was seeded onto the center of 3.5 cm slow-killing assay (SKA) plates, which were grown for 24 hours at 37°C and for another 24 hours at room temperature. Forty synchronized young adult animals were washed off of OP50 plates, washed three times in M9 buffer, and transferred to the assay plates (0.5 cm off the edge of the bacterial lawn), incubated at 25°C, and scored for avoidance 8 hours later.

Plates for the full-lawn killing assays were prepared essentially as above, with the exception that 20 μl of the overnight culture was spread to the edge of the plates. Forty synchronized young adult animals were added to the center of the lawns, incubated at 25°C, and scored for killing over the course of 7 days.

#### NO avoidance assay

Ten μl of an overnight OP50 culture was seeded at the center of 3.5 cm SKA plates, which were grown overnight at room temperature. NO donor solution was prepared freshly by dissolving DPTA NONOate (Cayman Chemical, #82110) in ddH2O to a final concentration of 100 mM. Forty synchronized young adult animals were washed off OP50 plates, washed three times in M9 buffer, and transferred to the assay plate (0.5 cm off the edge of the bacterial lawn). Immediately after, 10 ul of the NO donor solution was added on top of the OP50 lawn. The avoidance was scored 30 min later.

All assays were conducted by an experimenter unaware of genotype or experimental treatment. Four biological replicates were performed for each condition and genotype. Statistical analysis was performed as described in the figure legends.

### Fluorescence microscopy

Images were taken using an Olympus PlanAPO with a 100x 1.4 NA objective and an ORCA100 CCD camera (Hamamatsu). Young adult worms were immobilized with 30 mg/ml BDM (2,3-Butanedione monoxide, Sigma). Image stacks were captured and the maximum intensity projections were obtained using Metamorph 7.1 software (Molecular Devices).

### Calcium imaging

Calcium imaging was performed in a microfluidic device essentially as described (Chronis et al., 2007; Ha et al., 2010) with minor modification. Fluorescence time-lapse imaging was collected on a Nikon Eclipse Ti-U inverted microscope with a 40X oil immersion objective and a Yokogawa CSU-X1 scanner unit and a Photometrics CoolSnap EZ camera at 5 frames second^-1^. The GCaMP6.0s signal from the soma of the ASJ neurons was measured using NIH ImageJ. The change in the fluorescence intensity (ΔF) for each time point was the difference between its fluorescence intensity and the average intensity over the 20 seconds of the recording before stimulus onset (F_base_): ΔF = F-F_base_. Paired sample t-test was used for in-group statistical comparisons: for the response evoked by the onset of the NO donor, the average ΔF/F_base_% within a 3-second window prior to the stimulus onset was compared with the average ΔF/F_base_% within a 10-second window after the switch; for the response evoked by the removal of the NO donor, the average ΔF/F_base_% within a 3-second window prior to the stimulus removal was compared with the average ΔF/F_base_% within a 20-second window after the switch. Data was assessed for a normal distribution using the Shapiro-Wilk normality test. For comparisons of normally distributed data, a two-tailed Student’s t-test was used. For all other comparisons, a Wilcoxon signed rank test was used (SPSS Statistics). The NO donor solution was prepared freshly before recording by dissolving 10 mg of DPTA NONOate (Cayman Chemical, Item Number 82110) in 15 ml of nematode growth medium buffer (3 g/L NaCl, 1 mM CaCl2, 1 mM MgSO4, 25mM potassium phosphate buffer, pH6.0) and kept at room temperature for 30 minutes to allow adequate NO release before use.

## Acknowledgements

This work was supported by NIH Research Grants DK80215 (J.M.K.) and DC009852 (Y.Z.). We thank the *C. elegans Caenorhabditis* Genetic Center (supported by National Institutes of Health - Office of Research Infrastructure Programs (P40 OD010440) for strains, Cori Bargmann and Jihong Bai for construct(s), Constantine Haidaris and Fred Ausubel for *Pseudomonas aeruginosa* strains and protocols, and Antonio Miranda-Vizuete for strains and helpful discussions. We also thank members of the Kaplan and the Zhang laboratories, Fred Ausubel, and Josh Meisel for helpful suggestions, reagents and comments on this manuscript.

